# Selectivity to acoustic features of human speech in the auditory cortex of the mouse

**DOI:** 10.1101/2023.09.20.558699

**Authors:** Jennifer L. Mohn, Melissa M. Baese-Berk, Santiago Jaramillo

## Abstract

A better understanding of the neural mechanisms of speech processing can have a major impact in the development of strategies for language learning and in addressing disorders that affect speech comprehension. Technical limitations in research with human subjects hinder a comprehensive exploration of these processes, making animal models essential for advancing the characterization of how neural circuits make speech perception possible. Here, we investigated the mouse as a model organism for studying speech processing and explored whether distinct regions of the mouse auditory cortex are sensitive to specific acoustic features of speech. We found that mice can learn to categorize frequency-shifted human speech sounds based on differences in formant transitions (FT) and voice onset time (VOT). Moreover, neurons across various auditory cortical regions were selective to these speech features, with a higher proportion of speech-selective neurons in the dorso-posterior region. Last, many of these neurons displayed mixed-selectivity for both features, an attribute that was most common in dorsal regions of the auditory cortex. Our results demonstrate that the mouse serves as a valuable model for studying the detailed mechanisms of speech feature encoding and neural plasticity during speech-sound learning.

## 1 Introduction

Understanding the neural mechanisms underlying speech processing by the auditory system holds paramount importance in our comprehension of how we communicate, of the challenges of learning a new language, and of potential therapies for disorders of speech communication. Despite the wealth of knowledge gained from studies in human subjects (Yi et al., 2021; Oganian et al., 2023), the intricate neural processes involved in speech processing remain challenging to explore fully. Ethical considerations and technical limitations restrict the depth of insight that can be obtained solely through research in humans. A full understanding of how the brain processes and categorizes the acoustic features of speech requires descriptions of how neurons represent these features and how neural connections change when learning novel acoustic categories.

Because this level of investigation is extremely challenging to achieve in human subjects, an appropriate animal model is needed (Kluender, 2000; Lotto et al., 2003). Previous studies have demonstrated that various animal species are capable of learning phonetic categories that share perceptual qualities with humans (Kuhl and Padden, 1983; Kuhl and Miller, 1978; Engineer et al., 2015; Saunders and Wehr, 2019), suggesting that animal models are appropriate for the study of human speech processing by the auditory system. In particular, the mouse provides an unparalleled level of experimental access to investigate how the activity of individual neurons represent distinct acoustic features and how the circuits these neurons form change during the learning process. However, acoustic communication in mice is markedly different from that in humans. Mouse vocalizations are ultrasonic, the repertoire of “syllables” is more limited, and there is evidence that mouse vocalization is not learned, as vocalizations emitted by deaf mice do not differ in either structure or usage from hearing mice (Portfors, 2007; Portfors and Perkel, 2014; Hammerschmidt et al., 2012; Mahrt et al., 2013). Here, we assessed whether the mouse can serve as a good model organism for studying the neural mechanisms of speech processing, and evaluated whether neurons in distinct regions of the mouse auditory cortex are sensitive to specific features of speech.

We report that mice are able to learn to categorize frequency-shifted human speech sounds that varied according to two features: (1) formant transitions (FT), the spectral change in formant frequencies during a consonant (*e.g.*, the difference between /ba/ and /da/); and (2) voice onset time (VOT), the period of time from the burst of a plosive to the onset of the vocal fold vibration (*e.g.*, the difference between /ba/ and /pa/). Moreover, we show that neurons across many auditory cortical areas of the mouse (including primary, dorsal, and ventral auditory cortex, as well as temporal association area) are selective to these acoustic features, with the dorso-posterior region of the auditory cortex containing a higher proportion of speech-selective neurons compared to other areas. These combined behavioral and physiological results demonstrate that the mouse can serve as a valid model organism for investigating the encoding of speech features by the auditory system and how these neural representations may change throughout learning.

## 2 Materials and Methods

### 2.1 Animals

A total of 25 adult C57BL/6J mice (JAX 000664) of both sexes were used in this study. All procedures followed the National Institutes of Health animal care standards and were approved by the University of Oregon Animal Care and Use Committee. Animals were housed in group cages unless implanted for recording, in which case they were individually housed after surgery to prevent damage to implants. Mice used for electrophysiology were between 8 and 18 weeks old at the time of implantation. Mice used for behavioral experiments were between 12 and 13 weeks old at the start of behavioral training. All mice were maintained on a 12h light/dark cycle.

### 2.2 Auditory stimuli

Speech sounds were generated using Praat (www.praat.org) and the Python wrapper Parselmouth (Jadoul et al., 2018). Consonant-Vowel (CV) syllables (240 ms long) with different Formant Transitions (FT) and Voice Onset Times (VOT) were generated at frequencies 8 times higher than that of normal human speech to better match the hearing range of mice. The vowel sound had formants at F0=800 Hz, F1=5680 Hz, F2=9920 Hz, and F3=20000 Hz, corresponding to the vowel /a/. VOT sounds were generated by changing the voicing point from 2 ms to 64 ms (logarithmically spaced). The FT sounds were generated by changing the slopes of formants F2 and F3 from +9.1 to -9.1 octaves/s for the human-resembling sounds, or from -28.6 to 28.6 octaves/s for the mouse vocalization-resembling sounds. The average intensity of sounds was calibrated to be around 60 dB-SPL.

In addition to speech sounds, we presented simpler sounds during electrophysiological recordings. One set consisted of 100 ms pure tones (60 and 70 dB-SPL) of 16 different frequencies from 2 kHz to 40 kHz (logarithmically spaced) presented with an interstimulus interval randomized in the range 1–1.4 seconds. We presented 20 repetitions per frequency, randomly sorted. A different set of sounds consisted of sinusoidally amplitude modulated white noise at 11 modulation rates logarithmically spaced between 4 and 128 Hz (100% modulation depth, 500 ms duration, 60 dB-SPL max, 20 repetitions per condition, randomly sorted).

During electrophysiological recordings, auditory stimuli were presented in open-field configuration from a speaker (MF1, Tucker-Davis Technologies) contralateral to the side of recording.

### 2.3 Behavioral training and assessment

The two-alternative choice sound categorization task was carried out inside single-walled sound-isolation boxes (IAC-Acoustics). Behavioral data was collected using the *taskontrol* platform developed in our laboratory (www.github.com/sjara/taskontrol) using the Python programming language. Mice initiated each trial by poking their noses into the center port of a three-port behavior chamber. After a silent delay of random duration (150–250 ms, uniformly distributed), a speech sound was presented. Animals were required to choose one of the two side ports to obtain reward (2 *μ*l of water) according to the identity of sound (*e.g.*, left port for /ba/ and right port for /pa/). If animals chose the incorrect side port, a new trial had to be initiated.

To evaluate psychometric performance for different values of VOT or FT, we synthesized 6 sounds along each feature variation, with 3 sounds indicating reward on the left side and the other 3 sounds indicating reward on the right. The sound on each trial was randomized. During behavioral assessment, a particular cohort of mice was only exposed to variations in either VOT or FT.

Behavioral training consisted of a sequence of stages where animals were first familiarized with the reward ports. They were then required to initiate trials by poking in the center port (obtaining reward after correct choices even if they had made an incorrect choice first), before moving to the stage where they were required to choose the appropriate side port for reward on the first try after each stimulus. During these training stages, animals were presented with only the most extreme values of either VOT and FT, and only after achieving 70% correct for each sound within a session were the sounds with intermediate feature values added to evaluate psychometric performance.

### 2.4 Surgery

To prepare for head-fixed recordings, mice were implanted with a headbar and bilateral craniotomies were made above auditory cortex. Mice were anesthetized with isoflurane and placed into a stereotactic apparatus (Kopf instruments), a portion of the scalp was then removed and craniotomies were made bilaterally above auditory cortex. The dura mater was removed. A plastic well was fitted and attached to the skull surrounding each craniotomy. Each well was then filled with a silicone elastomer (Sylgard 170, Dow-Corning) to protect the brain. Mice were allowed to recover for at least 3 days before beginning electrophysiological recordings.

### 2.5 Electrophysiological recordings

Electrophysiological sessions were conducted inside single-walled sound-isolation boxes (IAC Acoustics, Naperville, IL). Electrical signals were collected using Neuropixels 1.0 probes (IMEC) via an NI PXIe-8381 acquisition module and the OpenEphys software (www.open-ephys.org). Probes were coated with a fluorescent dye (DiI: Cat# V22885, or DiD Cat# V22887, Thermo-Fisher Scientific) before penetration of the brain to allow for identification of the recording locations post-mortem. During the experiment, mice were head-fixed and allowed to run atop a wheel. Once the mouse was situated, the silicone elastomer above one of the craniotomies was removed, and the Neuropixels probe was lowered vertically into the auditory cortex with the probe tip reaching approximately 3 mm below the brain surface.

### 2.6 Histology and definition of brain areas

At the conclusion of the experiments, animals were euthanized with euthasol and perfused through the heart with 4% paraformaldehyde. Brains were extracted and left in 4% paraformaldehyde for at least 24 hours before slicing. Brain slices (thickness 100 *μ*m) were prepared under phosphate-buffered saline using a vibratome (Leica VT1000 S) and imaged using a fluorescence microscope (Axio Imager 2, Carl Zeiss) with a 2.5x objective. To determine the location of our recording sites, we manually registered each brain slice to the corresponding coronal section in Allen Mouse Brain Common Coordinate Framework (CCF) (Wang et al., 2020). From this registration, we estimated the location of each recorded cell in CCF coordinates. For our analysis, we combined the Dorsal Auditory Area and the Posterior Auditory Area into a single area we refer to as Dorsal Auditory Area (AudD).

In addition to parcellating our data according to atlas-defined auditory areas, we also parcellated the recordings according to dorso-ventral (D-V) and anterior-posterior (A-P) coordinates, independent of the boundaries between atlas-defined auditory cortical areas. To do this, we estimated the most anterior, posterior, dorsal, and ventral points of the auditory areas and temporal association area from the Allen CCF Atlas. We used these points to create a rectangle surrounding the auditory areas, and divided this rectangle into equal quadrants: DP, DA, VP and VA.

### 2.7 Analysis of neural data

Spiking activity of single units was isolated using the spike sorting package Kilosort (Pachitariu et al., 2016), available at https://github.com/MouseLand/Kilosort, and manually curated using Phy (Rossant et al., 2016), available at https://github.com/cortex-lab/phy/. The resulting data were analyzed using in-house software developed in Python (www.python.org).

Responsiveness to sounds was evaluated by calculating whether there was a statistically significant difference between evoked firing (throughout the duration of each sound) and baseline activity (-200 ms to 0 ms from sound onset), for any of the sounds presented, using a Wilcoxon signed-rank test with Bonferroni correction for multiple-comparisons. For pure-tone selectivity, a Gaussian function was fit to the spike counts across frequencies and the best frequency was defined as the peak of this Gaussian function. For evaluating if cells were significantly responsive to tones within the F2–F3 frequency range (7.1 kHz – 25 kHz), we estimated the cells’ responsiveness to each tone (Wilcoxon signed-rank test with p < Bonferroni corrected *α* = 0.05/16 = 0.003), and a cell was categorized as responsive in the F2–F3 range if it was significantly responsive to any of the tones within that frequency range.

Selectivity to each feature of speech was evaluated for each cell by calculating a selectivity index 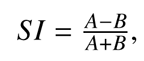 where A is the average firing rate from the stimulus that evoked the maximum change from baseline firing rate, and B is the average firing rate from the stimulus that evoked the minimum change from the baseline firing rate. For each SI calculation, we held the other (irrelevant) feature constant (*e.g.*, using the four stimuli represented on the bottom row of Fig. 2C). We chose to hold the irrelevant feature at either the maximum or the minimum value by assessing the condition in which the greatest change was observed between baseline and stimulus-evoked firing rates. To assess if the selectivity of a neuron to a specific feature was statistically significant, we performed a permutation test with 2000 repetitions, where the stimulus associated with each trial was randomized. A cell was categorized as selective for a given feature if the permutation test p-value was less than 0.05. A cell was determined to exhibit mixed-selectivity if it was significantly selective to both VOT and FT.

## 3 Results

### 3.1 Mice can discriminate features of speech

To test whether the mouse could be an appropriate animal model for studying the discrimination of speech sound features, we first tested whether mice are capable of learning to categorize human speech sounds. We generated consonant-vowel human speech sounds that were frequency shifted into the mouse hearing range (Fig. 1B). These sounds varied in one of two features: formant transitions (FT), the spectral change in formant features during a consonant (*e.g.*, the difference between /ba/ and /da/); or voice onset time (VOT), the period of time from the burst of a plosive to the onset of the vocal fold vibration (*e.g.*, the difference between /ba/ and /pa/). We trained mice to categorize speech sounds in a two-alternative choice task (Fig. 1A), based either on changes in VOT or FT, starting with extreme values of these features and then adding sounds with intermediate feature values to characterize the psychometric performance for each feature variation.

**Figure 1:**
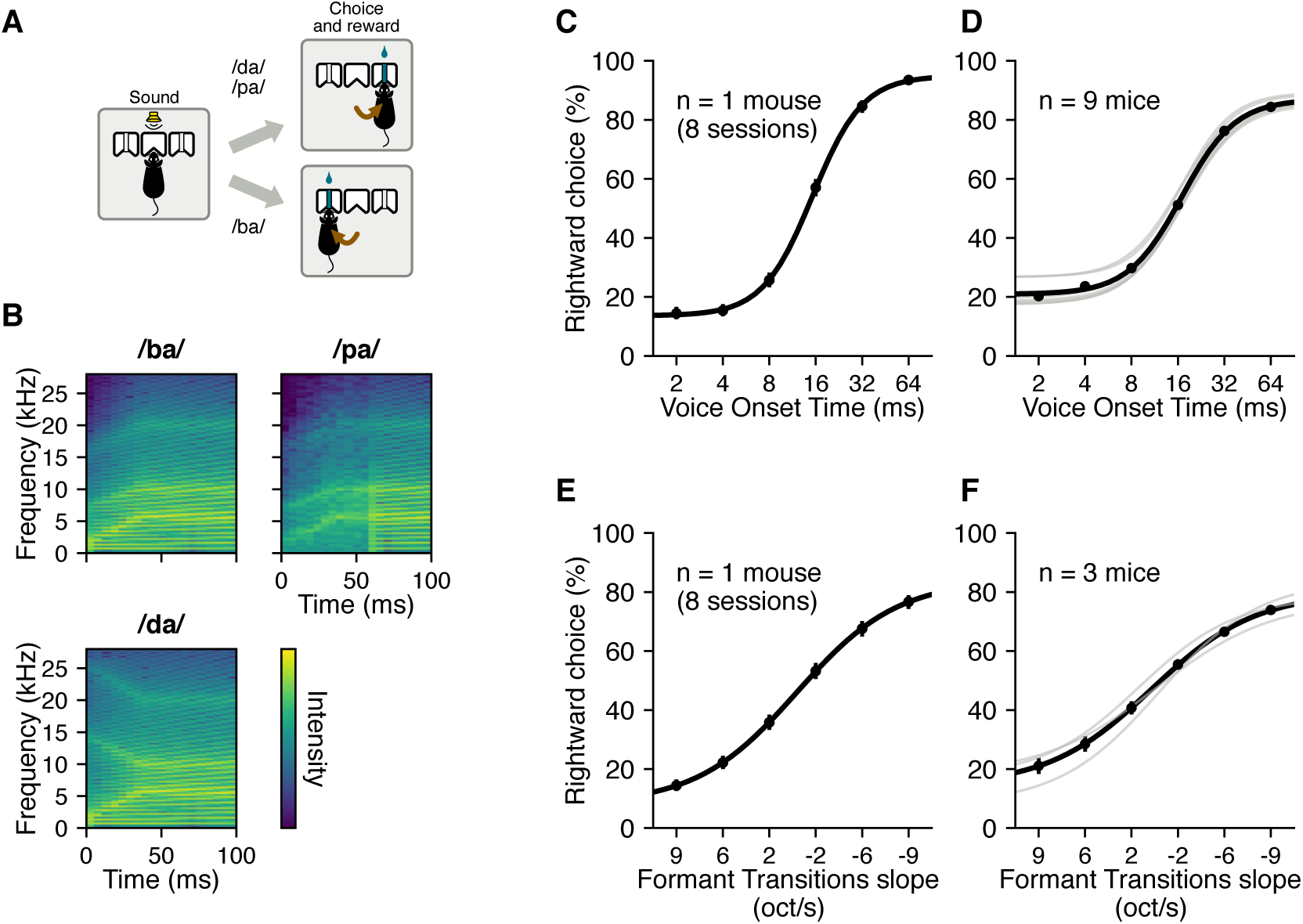
Mice can categorize speech sounds. **(A)** Two-alternative choice speech categorization task. Mice initiated a trial by entering the center port, triggering a sound presentation. They then had to choose the left or right port based on which sound was presented: left for short VOT and upwards sloping FT (/ba/), right for long VOT (/pa/), or downwards sloping FT, (/da/). A correct choice resulted in a water reward, an incorrect choice ended the trial without reward. **(B)** Spectrograms of /ba/, /da/, and /pa/ sounds presented (shifted to the mouse hearing range). Sounds were 240 ms long (only the first 100 ms shown here). Sounds with VOTs intermediate to /ba/ and /pa/ and FTs intermediate to /ba/ and /da/ were also presented. **(C)** Psychometric curve from a single mouse trained to categorize sounds by VOT. Error bars indicate 95% confidence interval. **(D)** Average psychometric curve from the VOT cohort (black) and psychometric fit for each mouse (gray). Error bars indicate SEM across mice. **(E)** As in C, for a mouse trained to categorize sounds by FT. **(F)** As in D, for the cohort trained to categorize FT.

Mice were able to successfully categorize sounds according to both VOT (Fig. 1C,D) and FT (Fig. 1E,F), as illustrated by the approximately 80% correct on the extremes of both VOT value and FT slope. As expected, intermediate values of the speech features resulted in performance closer to chance level (50%). While all of the VOT animals successfully categorized sounds by VOT (9 out of 9 total mice), fewer of the FT animals were able to successfully categorize sounds by FT throughout the training period (3 out of 9 total mice, 106 training sessions). It took on average 49 sessions (min = 30 sessions, max = 73 sessions) for the animals trained to categorize VOT to achieve 70% correct performance categorizing the two extremes of VOT sounds presented (VOT = 2ms and VOT = 64ms). The animals trained to categorize FT took longer to succeed at categorizing the extreme FT sounds presented (+/-9.1 oct/s), on average 64 sessions (min = 56 sessions, max = 76 sessions).

Since the mice were less successful at learning to categorize FT sounds compared to VOT sounds, we tested whether using sounds with FT slopes that more closely resemble the frequency sweeps that occur in mouse vocalizations would result in better performance. To do this, we generated a new set of FT sounds, such that the extreme slopes were +/-28.6 oct/s, which more closely resemble the features of mouse vocalizations (Portfors, 2007), compared to the +/-9.1 oct/s of the human-resembling FT sounds. We trained a new cohort of mice to discriminate these new FT sounds. A larger fraction of mice in this new cohort was able to learn to categorize FT extremes (8 out of 9 total mice), and they reached 70% correct performance more quickly in an average of 48 sessions (min = 26 sessions, max = 67 sessions). Overall, these results suggest that while the categorization performance of speech features depends on the specific FT slopes used, mice can learn to discriminate basic features of human speech sounds.

### 3.2 Neurons across the auditory cortex of mice are responsive to speech sounds

After verifying that mice were able to perceive differences in acoustic features of speech, we wanted to characterize neural responses to these sounds across auditory cortical areas of the mouse. We recorded extracellular electrophysiological responses of neurons from primary and non-primary auditory cortex, as well as the temporal association cortex (TeA) using Neuropixels 1.0 probes (Fig. 2A, 2B). We presented 12 consonant-vowel stimuli that varied in both FT and VOT (Fig. 2C) to awake, naive, head-fixed mice (n = 7). Additionally, prior to presenting speech sounds, we presented pure tones of varying frequencies and amplitude modulated noise (AM) of varying rates during each recording session in order to characterize the responses of neurons to simpler sound features. We recorded a total of 1009 single neurons from the auditory cortex and surrounding areas. Of these, 563 (55.8%) were responsive to any of the sounds presented (Fig. 2D, red + blue), and 347 cells were responsive to any of the speech sounds we presented (34.3% of all cells, 61.6% of sound responsive cells; Fig. 2D, red). A cell was classified as responsive to a class of sounds presented if it had a statistically significant difference between the evoked firing rate and baseline firing rate (Wilcoxon signed-rank test, p < 0.05, Bonferroni corrected alpha within each class of sounds given the number of stimuli presented).

**Figure 2:**
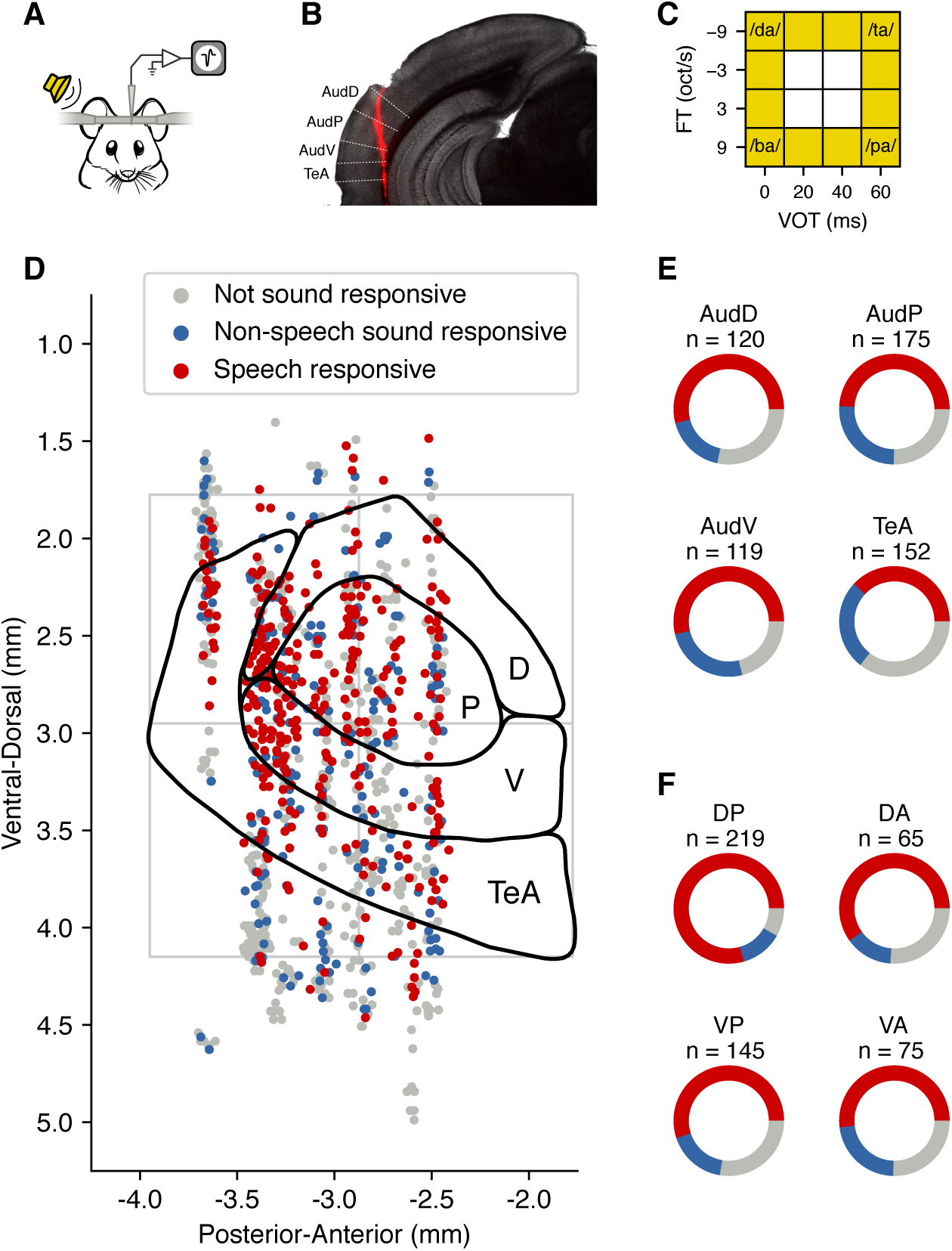
Electrophysiological recordings from multiple auditory cortical areas. **(A)** Single-neuron extracellular electrophysiology was recorded from awake, head-fixed mice (n = 7) using Neuropixels 1.0 probes. Mice naive to the behavioral task listened passively while sounds were presented contralaterally to the recording site. **(B)** Example electrode penetration spanning multiple areas of the auditory cortex, shown on a coronal slice. **(C)** Sounds presented during the recorded sessions varied in FT and VOT. Colored squares indicate presented sounds, empty squares indicate combinations that were not presented. **(D)** Locations of all cortical cells recorded across all mice, colored according to their responsiveness to sounds. Approximate boundaries of auditory brain areas according to the Allen Mouse Brain Atlas. P: primary, D: dorsal (combining AudPo and AudD), V: ventral, TeA: temporal association area. Gray lines show the borders of each quadrant used for the analysis in F. **(E)** The proportion of speech responsive cells was similar between each of the atlas-defined areas. **(F)** Parcellating the auditory cortex into regions along the A-P/D-V axes yielded a higher proportion of speech responsive cells in the dorso-posterior region.

Once we knew there were cells that were responsive to these speech sounds, we tested whether the proportions of cells that were speech responsive were different across the cortical areas recorded. Using functional and anatomical mapping of the mouse auditory cortex, previous studies have parcellated mouse auditory cortex into anywhere from 3 to 6 sub-areas (Ceballo et al., 2019; Romero et al., 2020; Issa et al., 2017; Tsukano et al., 2016). Because of these discrepancies, we chose to follow two distinct approaches for investigating differences in neural responses across auditory cortical regions. We first evaluated neural activity according to cortical areas defined in the Allen Mouse Brain Atlas: primary auditory cortical area (AudP), dorsal and posterior auditory areas (combined and referred to here as AudD), ventral auditory area (AudV) and TeA, after registering each mouse brain to the atlas. We also used a different approach where we parcellated the whole auditory cortical region according to dorso-ventral (D-V) and anterior-posterior (A-P) coordinates, independent of the boundaries between atlas-defined auditory cortical areas.

When we compared the fraction of speech responsive cells (Fig. 2E, red) as a proportion of total cells (Fig. 2E, all cells) across each atlas-defined area, we found that there were no statistically significant differences in the fraction of speech responsive cells between any of these areas (Table 1). Further, when we restricted our analysis to sound responsive cells (Fig. 2E, red + blue), the proportion of cells that were responsive to the speech sounds (Fig. 2E, red) was also similar across auditory cortical areas (Table 2).

**Table 1:**
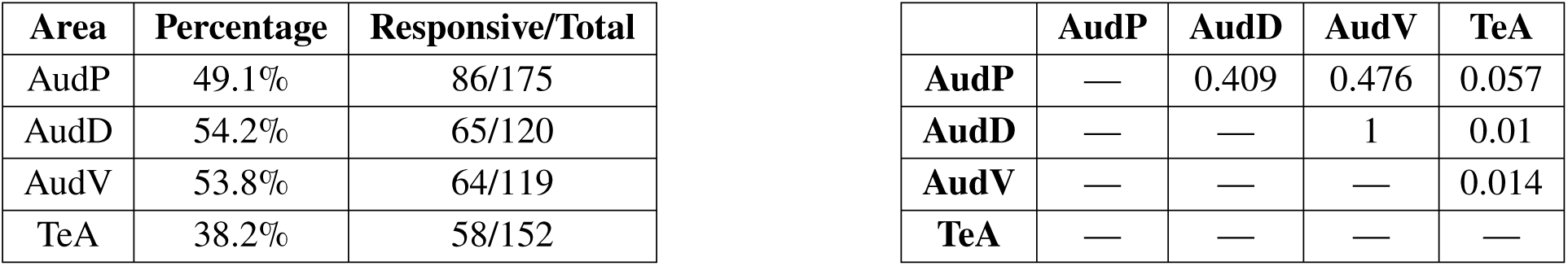
The proportion of speech responsive cells with respect to total cells recorded is similar across auditory areas. Right: p-values from comparisons between areas (Fisher Exact Test, Bonferroni corrected *α* = 0.05/6 = 0.008).

Tables associated with Figure 2
| Area | Percentage | Responsive/Total |
| --- | --- | --- |
| AudP | 49.1% | 86/175 |
| AudD | 54.2% | 65/120 |
| AudV | 53.8% | 64/119 |
| TeA | 38.2% | 58/152 |

**Table 1:** The proportion of speech responsive cells with respect to total cells recorded is similar across auditory areas. Right: p-values from comparisons between areas (Fisher Exact Test, Bonferroni corrected $\alpha = 0.05/6 = 0.008$ ).
|  | AudP | AudD | AudV | TeA |
| --- | --- | --- | --- | --- |
| **AudP** | — | 0.409 | 0.476 | 0.057 |
| **AudD** | — | — | 1 | 0.01 |
| **AudV** | — | — | — | 0.014 |
| **TeA** | — | — | — | — |

**Table 2:** The proportion of speech responsive cells with respect to sound responsive cells is similar across auditory areas. Right: p-values from comparisons between areas (Fisher Exact Test, Bonferroni corrected *α* = 0.05/6 = 0.008).

| Area | Percentage | Speech resp/<br>Sound resp |
| --- | --- | --- |
| AudP | 65.6% | 86/131 |
| AudD | 75.6% | 65/86 |
| AudV | 68.1% | 64/94 |
| TeA | 59.2% | 58/98 |

**Table 2:** The proportion of speech responsive cells with respect to sound responsive cells is similar across auditory areas. Right: p-values from comparisons between areas (Fisher Exact Test, Bonferroni corrected $\alpha = 0.05/6 = 0.008$ ).
|  | AudP | AudD | AudV | TeA |
| --- | --- | --- | --- | --- |
| **AudP** | — | 0.133 | 0.775 | 0.336 |
| **AudD** | — | — | 0.321 | 0.02 |
| **AudV** | — | — | — | 0.231 |
| **TeA** | — | — | — | — |

In contrast, when we parcellated our data by quadrants in the A-P and D-V directions, instead of atlas-defined areas, we found differences in the proportion of speech responsive cells (Fig. 2F, red) between cortical regions. As a proportion of total cells in each quadrant (Fig. 2F, all cells), we found that the dorso-posterior region had significantly more speech responsive cells (79.5%) compared to each of the other regions (60.0% DA, 55.2% VA, 52.0% VP, Table 3). Further, when we restricted our analysis to the sound responsive cells (Fig. 2F, red + blue) we found that the DP region still had a higher proportion of speech responsive cells compared to VA, but not DP or VP (Table 4). In summary, while we observed no differences between the atlas-defined auditory cortical areas in the proportion of speech responsive cells out of either the total cells recorded or of sound responsive cells, neurons in the dorso-posterior region of the auditory cortex were more likely to be speech responsive compared to the ventral regions.

**Table 3:** The proportion of speech responsive cells with respect to all recorded cells is higher in the dorso-posterior region compared to the other regions. Right: p-values from comparisons between regions.

| Region | Percentage | Responsive/Total |
| --- | --- | --- |
| DP | 79.5% | 174/219 |
| DA | 60.0% | 39/65 |
| VP | 55.2% | 80/145 |
| VA | 52.0% | 39/75 |

|  | DP | DA | VP | VA |
| --- | --- | --- | --- | --- |
| **DP** | — | 0.003 | <0.001 | <0.001 |
| **DA** | — | — | 0.549 | 0.395 |
| **VP** | — | — | — | 0.671 |
| **VA** | — | — | — | — |

**Table 4:** The proportion of speech responsive cells with respect to sound responsive cells is higher in the dorso-posterior region compared to the ventral-anterior region. Right: p-values from comparisons between regions (Fisher Exact Test, Bonferroni corrected *α* = 0.05/6 = 0.008).

| Region | Percentage | Speech resp/<br>Sound resp |
| --- | --- | --- |
| DP | 86.6% | 174/201 |
| DA | 81.2% | 39/48 |
| VP | 76.2% | 80/105 |
| VA | 69.6% | 39/56 |

**Table 4:** The proportion of speech responsive cells with respect to sound responsive cells is higher in the dorso-posterior region compared to the ventral-anterior region. Right: p-values from comparisons between regions (Fisher Exact Test, Bonferroni corrected $\alpha = 0.05/6 = 0.008$ ).
|  | DP | DA | VP | VA |
| --- | --- | --- | --- | --- |
| **DP** | — | 0.363 | 0.025 | 0.005 |
| **DA** | — | — | 0.536 | 0.256 |
| **VP** | — | — | — | 0.451 |
| **VA** | — | — | — | — |

### 3.3 Auditory cortical neurons change their firing rate in response to changing VOT and FT

Having established the presence of speech responsive cells in each of the regions we sampled, we wanted to test whether cells showed selectivity to features of speech or if they were indiscriminately responsive to the speech sounds we presented. We found some cells that changed their firing rate drastically when VOT changed, but had little change in their firing when FT changed (Fig. 3A). Conversely, there were cells that changed their firing rate drastically when FT changed, but their activity had little change when VOT changed (Fig. 3B). There were also cells whose firing rates varied depending on both FT and VOT (Fig. 3C). Finally, some cells were responsive to the speech sounds, but their responses did not change when varying either feature (Fig. 3D).

**Figure 3:**
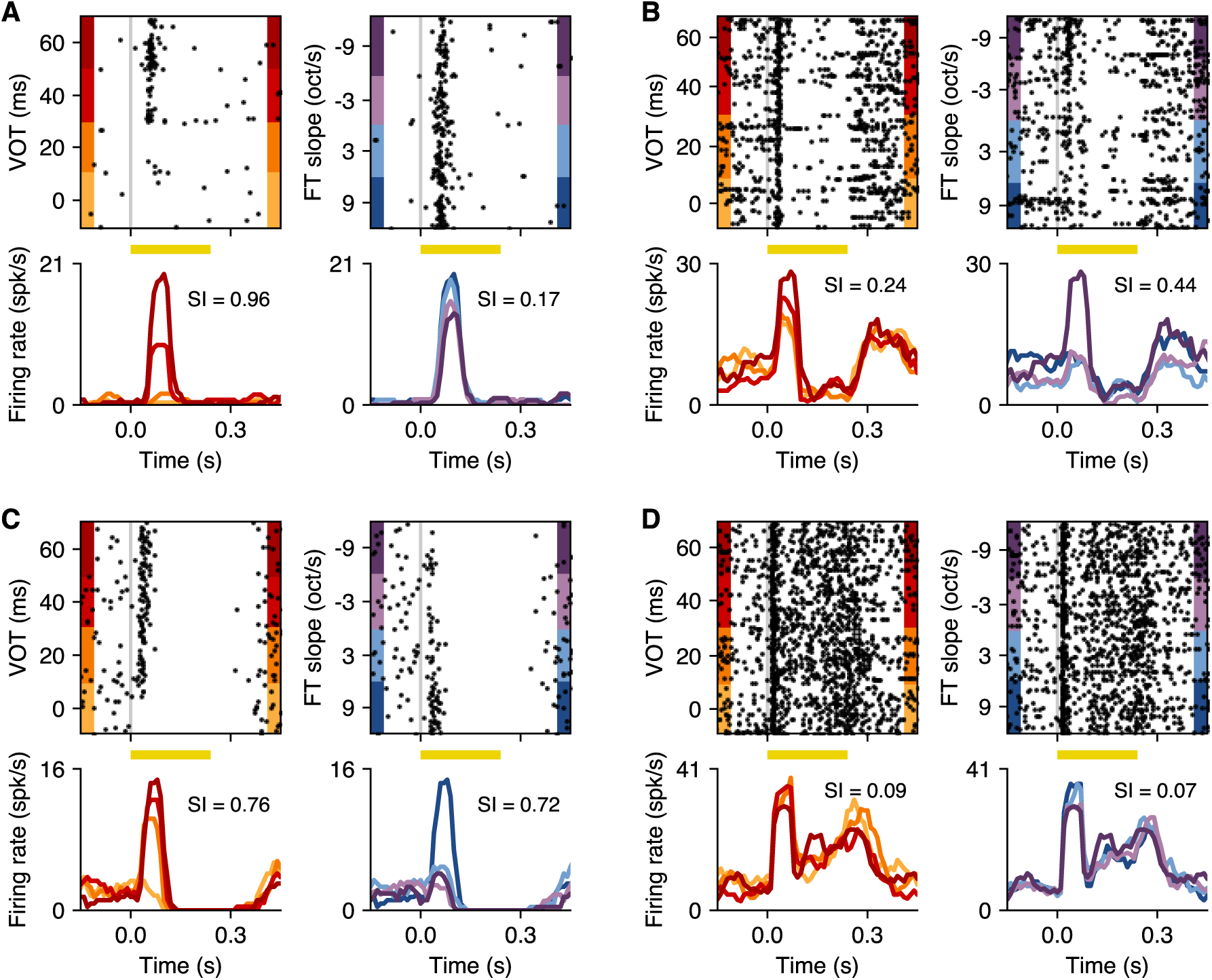
Neurons in the auditory cortex are selective to speech sounds. **(A)** Sound evoked responses from an example neuron that is significantly selective to VOT (left panel), but not to FT (right panel). The yellow horizontal bars represent when the auditory stimulus was presented. SI: selectivity index. **(B)** Example neuron that is significantly selective to FT, but not VOT. **(C)** Example neuron that is significantly selective to both VOT and FT, which we classify as mixed-selective. **(D)** Example neuron that is responsive to speech sounds, but not selective for either VOT or FT features.

To enable comparing selectivity across neurons and auditory cortical regions, we calculated for each neuron a selectivity index (SI) associated with each feature, while we held the other (irrelevant) feature constant (*e.g.*, using the four stimuli represented on the bottom row of Fig. 2C). We chose to hold the irrelevant feature at either the minimum or the maximum value by assessing the condition in which the greatest change was observed between baseline and stimulus-evoked firing rates. The SIs shown for each neuron in Fig. 3 support the qualitative observations described above illustrating the existence of cells selective to a single feature, cells that had mixed-selectivity (*i.e.*, selectivity to both features), and cells that were responsive to speech sounds but not selective to either feature.

### 3.4 Selectivity to speech features is greater in the dorso-posterior region of auditory cortex

To compare speech selectivity across areas of the auditory cortex, we first assessed whether the proportion of cells that were selective to each feature was different across the atlas-defined brain areas we sampled. All areas we tested contained cells that were selective to VOT (Fig. 4A). We found that of all the cells recorded, approximately 20-30% of cells in each area were selective to VOT (Fig. 4B), and there were no statistically significant differences in the proportion of cells that were selective to VOT between any of the areas (Table 5). There were also cells selective to FT in each of the areas we tested (Fig. 4E). Similar to VOT, there were no statistically significant differences in the proportion of FT selective cells between any of the areas we sampled (Table 6). Each area had between approximately 5-15% of cells that were FT selective (Fig. 4F). Similarly, when we assessed the proportion of cells selective to VOT or FT in each area with respect to the number of sound responsive cells or the number of speech responsive cells we found no statistically significant differences between any of the areas for either feature (Tables 5 & 6). In addition to calculating the proportion of cells that were selective to speech features in each area, we tested whether the strength of that selectivity was different between areas (Fig. 4A, 4E). When we compared the distribution of selectivity indices between areas among selective cells, we found that there was no difference in the distribution of VOT selectivity indices between areas (Kruskal-Wallis, p = 0.426). Likewise, there was no difference in the strength of FT selectivity between any of the areas we sampled (Kruskal-Wallis p= 0.907).

**Figure 4:**
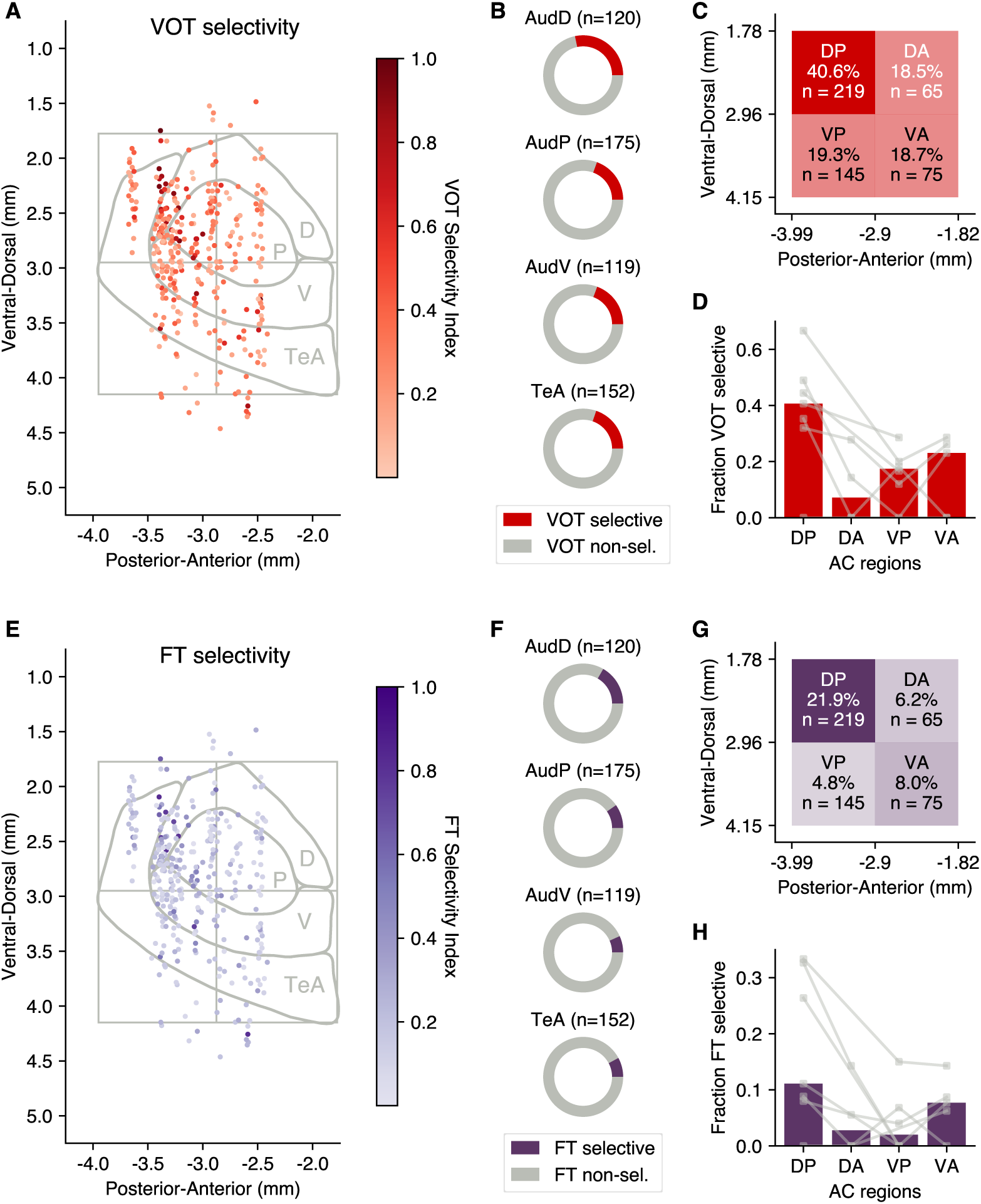
Selectivity to speech features is greater in the dorso-posterior region of auditory cortex. **(A)** Locations of speech responsive cells recorded, colored according to VOT selectivity indices. **(B)** The proportion of VOT selective cells is similar across atlas-defined areas. **(C)** The proportion of VOT selective cells is greater in the dorso-posterior region of auditory cortex. **(D)** Proportion of VOT selective cells in each region for each mouse. Dots and connecting lines indicate values for individual animals. Bars indicate medians across animals. Regions with fewer than 3 recorded cells were excluded. **(E-H)** As in A-D, for FT selectivity.

**Table 5:**
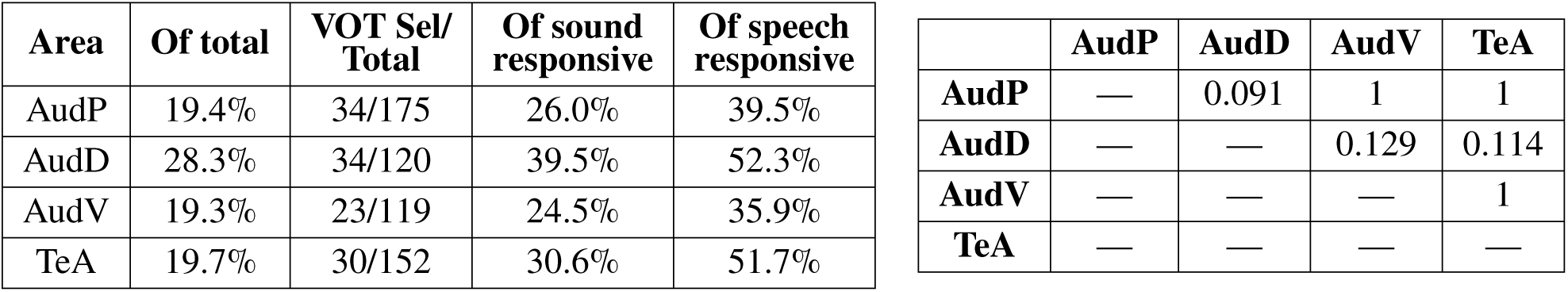
The proportion of cells selective to VOT out of total cells was similar between atlas-defined auditory areas. Right: p-values from comparisons between areas of VOT selective cells out of the total cells (Fisher Exact Test, Bonferroni corrected *α* = 0.05/6 = 0.008).

Tables associated with Figure 4
| Area | Of total | VOT Sel/<br>Total | Of sound<br>responsive | Of speech<br>responsive |
| --- | --- | --- | --- | --- |
| AudP | 19.4% | 34/175 | 26.0% | 39.5% |
| AudD | 28.3% | 34/120 | 39.5% | 52.3% |
| AudV | 19.3% | 23/119 | 24.5% | 35.9% |
| TeA | 19.7% | 30/152 | 30.6% | 51.7% |

**Table 5:** The proportion of cells selective to VOT out of total cells was similar between atlas-defined auditory areas. Right: p-values from comparisons between areas of VOT selective cells out of the total cells (Fisher Exact Test, Bonferroni corrected $\alpha = 0.05/6 = 0.008$ ).
|  | AudP | AudD | AudV | TeA |
| --- | --- | --- | --- | --- |
| **AudP** | — | 0.091 | 1 | 1 |
| **AudD** | — | — | 0.129 | 0.114 |
| **AudV** | — | — | — | 1 |
| **TeA** | — | — | — | — |

**Table 6:** The proportion of cells selective to FT out of total cells was similar between atlas-defined auditory areas. Right: p-values from comparisons between areas of FT selective cells out of the total cells (Fisher Exact Test, Bonferroni corrected *α* = 0.05/6 = 0.008).

| Area | Of total | FT Sel/<br>Total | Of sound<br>responsive | Of speech<br>responsive |
| --- | --- | --- | --- | --- |
| AudP | 9.7% | 17/175 | 13.0% | 19.8% |
| AudD | 16.7% | 20/120 | 23.3% | 30.8% |
| AudV | 6.7% | 8/119 | 8.5% | 12.5% |
| TeA | 7.9% | 12/152 | 12.2% | 20.7% |

**Table 6:** The proportion of cells selective to FT out of total cells was similar between atlas-defined auditory areas. Right: p-values from comparisons between areas of FT selective cells out of the total cells (Fisher Exact Test, Bonferroni corrected $\alpha = 0.05/6 = 0.008$ ).
|  | AudP | AudD | AudV | TeA |
| --- | --- | --- | --- | --- |
| **AudP** | — | 0.106 | 0.403 | 0.697 |
| **AudD** | — | — | 0.026 | 0.036 |
| **AudV** | — | — | — | 0.817 |
| **TeA** | — | — | — | — |

In contrast to the results from atlas-defined auditory cortical areas, when we parcellated our data into A-P/D-V quadrants we found clear differences in the selectivity to speech features between regions of the auditory cortex (Fig. 4C, 4G). We found that out of all cells, the dorso-posterior quadrant had a significantly higher proportion of cells selective to VOT compared to any of the other quadrants (DP: 40.6%, DA: 18.5%, VP: 19.3%, VA: 18.7%, Table 7). To verify that these results were not being driven by a single animal, we assessed the proportion of VOT selective cells for each region recorded in individual animals and found that this trend was common across most animals (Fig. 4D). Similarly, when we compared the proportion of FT selective cells between these regions, we found that the dorso-posterior region had a significantly higher proportion of FT selective cells compared to all the other regions (DP: 21.9%, DA: 6.2%, VP: 4.8%, VA: 8.0%), and found no statistically significant differences across any of the other regions (Table 8). To verify that this larger proportion of FT selective cells in DP compared to the other regions was not driven by a single animal, we assessed the proportion of FT selective cells in each region for individual animals and found that this trend was consistent across most animals (Fig. 4H). Lastly, the observation that the DP region had a higher proportion of VOT and FT selective cells compared to other regions was apparent also when we restricted our analysis to only sound responsive cells or speech responsive cells (Tables 7, 8), suggesting that this result is not simply the consequence of the cells’ responsiveness to the speech sounds presented.

**Table 7:** The proportion of cells selective to VOT out of total cells was greater in the dorso-posterior region of auditory cortex compared to the other regions. Right: p-values from comparisons between regions of VOT selective cells out of the total cells (Fisher Exact Test, Bonferroni corrected *α* = 0.05/6 = 0.008).

| Region | Of total | VOT Sel/<br>Total | Of sound<br>responsive | Of speech<br>responsive |
| --- | --- | --- | --- | --- |
| DP | 40.6% | 89/219 | 44.3% | 51.1% |
| DA | 18.5% | 12/65 | 25.0% | 30.8% |
| VP | 19.3% | 28/145 | 26.7% | 35.0% |
| VA | 18.7% | 14/75 | 25.0% | 35.9% |

**Table 7:** The proportion of cells selective to VOT out of total cells was greater in the dorso-posterior region of auditory cortex compared to the other regions.
|  | DP | DA | VP | VA |
| --- | --- | --- | --- | --- |
| **DP** | — | 0.001 | <0.001 | <0.001 |
| **DA** | — | — | 1 | 1 |
| **VP** | — | — | — | 1 |
| **VA** | — | — | — | — |

**Table 8:**
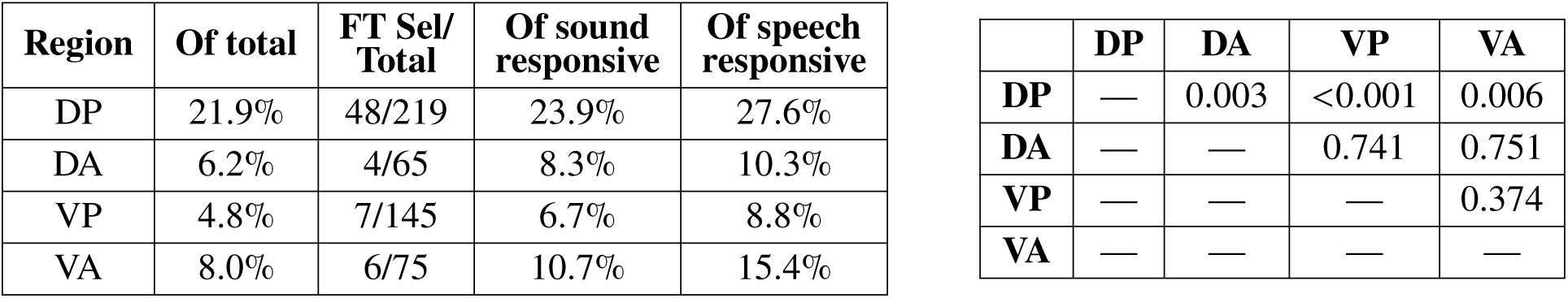
The proportion of cells selective to FT out of total cells was greater in the dorso-posterior region of auditory cortex compared to the other regions. Right: p-values from comparisons between regions FT selective cells out of the total cells (Fisher Exact Test, Bonferroni corrected *α* = 0.05/6 = 0.008).

In summary, while we observed no differences in selectivity to speech features between atlas-defined auditory cortical areas, when we parcellated our recorded region into quadrants, we found that neurons in the dorso-posterior region of auditory cortex were more likely to be selective to both VOT and FT compared to the other regions of auditory cortex.

### 3.5 Selectivity to speech features is not explained by frequency selectivity

We next wanted to test whether the differences we found in selectivity to speech features across regions could be explained simply by differences in selectivity to sound frequency. In particular, since FT sounds are composed of varying frequencies in the F2 and F3 formants (7.1kHz – 25kHz in the sounds presented, Fig. 1B), it is possible that the differences we observed across regions in selectivity to FT could be explained simply by differences in frequency selectivity given the sounds we presented, rather than selectivity to the speech feature. We focused on FT and not VOT selectivity, because the changes in VOT sounds occur across all of the frequencies, and therefore one does not expect frequency selectivity to explain differences in VOT selectivity. To test whether selectivity to FT may depend on frequency selectivity, we first calculated the best frequency (BF) for each sound responsive cell and compared the distribution of BFs between regions. We found that there was a statistically significant difference in the median best frequency of cells between quadrants (p<0.001, Kruskal-Wallis), with median values falling within the F2–F3 range: 11.1 kHz for DP, 23.1 kHz for DA, 10.3 kHz for VP and 24.3 kHz for VA.

However, one would expect that any neuron that responds to tones within the F2–F3 frequency range could be selective to FT, even if their best frequency is not in this range. Therefore, we focused on cells that had a statistically significant response to pure tone frequencies in the F2–F3 range. When we restrict our analysis of FT selectivity to these subsets of cells, we found that DP had 29.4% of cells selective to FT, while DA only had 9.1%, VP had 10.7%, and VA had 17.2%. Even though in this case only differences between DP and VP reached statistical significance (Table 9), these trends are consistent with the results we found using the whole data set. These result suggests that the differences in FT selectivity we found between regions cannot be fully explained by distinct selectivity to the frequency of pure tones.

**Table 9:**
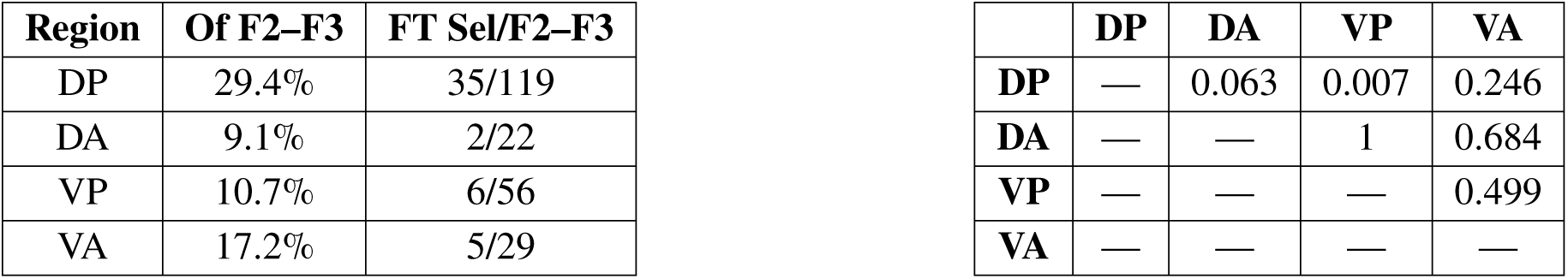
The proportion of cells selective to FT out of cells responsive to tones in the F2–F3 frequency range was significantly higher in the dorso-posterior region of auditory cortex compared to the ventro-posterior region. Right: p-values from comparisons between regions (Fisher Exact Test, Bonferroni corrected *α* = 0.05/6 = 0.008).

### 3.6 Mixed-selectivity is more prevalent in the dorsal portion of auditory cortex

In addition to cells that were selective to just VOT or FT, we found cells that changed their firing rate in response to both features (Fig. 3C). We defined a cell as having mixed-selectivity if it was significantly selective to both VOT and FT. If the cell was significantly selective to only one of the two features, it was classified as single-feature selective. Of the speech-selective cells, we found cells with mixed selectivity in each of the areas we examined (Fig. 5A). We then wanted to test if the distribution of mixed-selective cells was different across auditory cortical areas. When we parcellated our data by atlas-defined cortical areas, we found that out of the speech responsive cells, there were significantly fewer mixed-selective cells in AudV (6.2%) compared to AudD (24.6%, Table 10, Fig. 5B). While the proportion of mixed-selective cells was trending lower in AudV compared to both AudP and TeA, there was no statistically significant difference in the proportion of mixed-selective cells out of speech responsive cells among these areas (Table 10). Restricting the analysis to only speech-selective cells, yielded similar trends, although there were no statistically significant differences between the areas.

**Figure 5:**
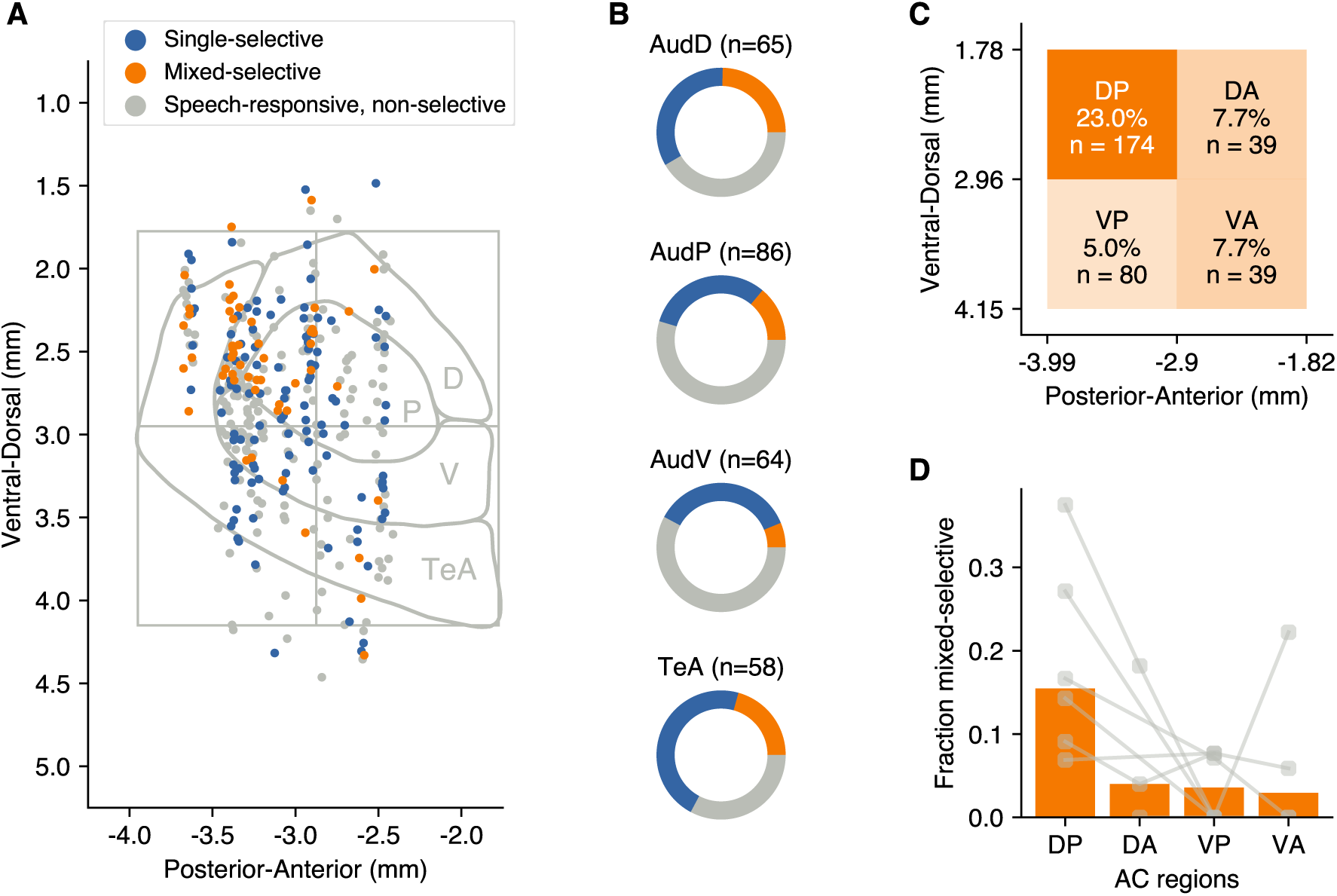
Mixed selectivity is more prevalent in the dorsal auditory areas. **(A)** Locations of speech responsive cells recorded, colored according to their selectivity to speech features. **(B)** The proportion of mixed-selective cells is higher in AudD compared to AudV. **(C)** The proportion of mixed-selective cells is 3-5 times as high in the DP region compared to the other auditory cortical regions. **(D)** Proportion of mixed-selective cells out of speech responsive cells for individual animals. Dots and connecting lines indicate values for individual animals. Instances in which an individual had fewer than 3 recorded cells in a region were excluded. Bars indicate the median proportion for each region averaged across animals.

**Table 10:**
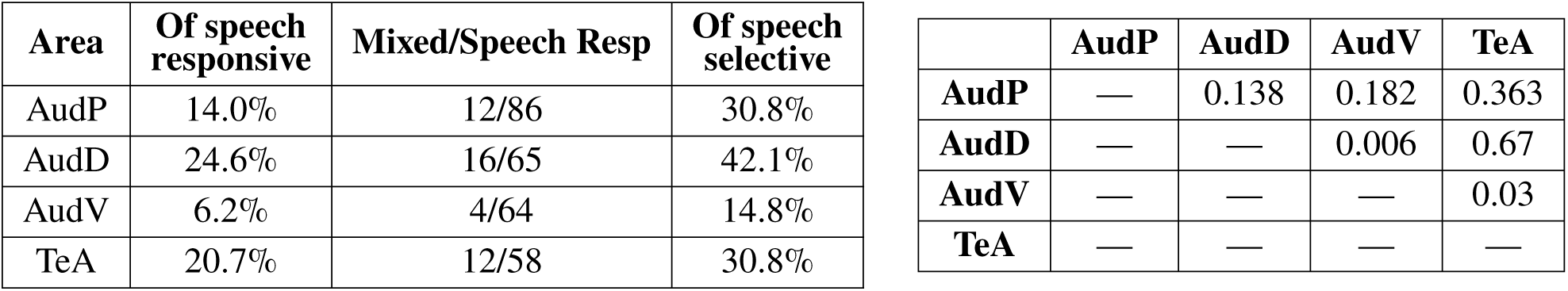
The proportion of cells with mixed-selectivity out of speech responsive cells was higher in dorsal auditory area compared to the ventral auditory area. Right: p-values from comparisons between areas of mixed-selective cells out of the speech responsive cells (Fisher Exact Test, Bonferroni corrected *α* = 0.05/6 = 0.008).

Tables associated with Figure 5
| Area | Of speech responsive | Mixed/Speech Resp | Of speech selective |
| --- | --- | --- | --- |
| AudP | 14.0% | 12/86 | 30.8% |
| AudD | 24.6% | 16/65 | 42.1% |
| AudV | 6.2% | 4/64 | 14.8% |
| TeA | 20.7% | 12/58 | 30.8% |

**Table 10:** The proportion of cells with mixed-selectivity out of speech responsive cells was higher in dorsal auditory area compared to the ventral auditory area.
|  | AudP | AudD | AudV | TeA |
| --- | --- | --- | --- | --- |
| AudP | — | 0.138 | 0.182 | 0.363 |
| AudD | — | — | 0.006 | 0.67 |
| AudV | — | — | — | 0.03 |
| TeA | — | — | — | — |

As before, we wanted to test whether there were any differences in the proportion of mixed-selective cells across the A-P/D-V axes, by parcellating the auditory cortex region into quadrants (Fig. 5C). We found that the dorso-posterior quadrant had over twice the proportion of mixed-selective cells compared to each of the other areas (DP: 23.4%, DA: 7.7%, VP: 5.0%, VA: 7.7%), although only the difference between regions DP and VP was statistically significant (Table 11). This trend, where region DP had a higher proportion of mixed-selective cells was consistent across individual animals (Fig. 5D). In summary, we found there to be a higher proportion of mixed-selective cells in the atlas-defined dorsal auditory cortex compared to the ventral area, and when parcellating the auditory cortex into quadrants the results followed a similar trend with the dorso-posterior region having a higher proportion of mixed-selective cells than the ventro-posterior region.

**Table 11:** The proportion of cells with mixed-selectivity out of speech responsive cells was higher in dorso-posterior region compared to the ventral-posterior region. Right: p-values from comparisons between regions of mixed-selective cells out of the speech responsive cells (Fisher Exact Test, Bonferroni corrected *α* = 0.05/6 = 0.008).

| Region | Of speech responsive | Mixed/Speech Resp | Of speech selective |
| --- | --- | --- | --- |
| DP | 23.0% | 40/174 | 41.2% |
| DA | 7.7% | 3/39 | 23.1% |
| VP | 5.0% | 4/80 | 12.9% |
| VA | 7.7% | 3/39 | 17.6% |

**Table 11:** The proportion of cells with mixed-selectivity out of speech responsive cells was higher in dorso-posterior region compared to the ventro-posterior region.
|  | DP | DA | VP | VA |
| --- | --- | --- | --- | --- |
| DP | — | 0.045 | <0.001 | 0.045 |
| DA | — | — | 0.682 | 1 |
| VP | — | — | — | 0.682 |
| VA | — | — | — | — |

## 4 Discussion

Our behavioral results demonstrate that mice are able to discriminate changes in key acoustic features of human speech, namely, formant transitions and voice onset time. Our finding extend recent results showing that mice can learn to discriminate speech sounds like “sad” vs. “dad” (O’Sullivan et al., 2020) or categorize consonant-vowel pairs for consonants /g/ and /b/ and generalize these categories after changing the speaker or vowel context (Saunders and Wehr, 2019). Our study also demonstrates that subregions of the auditory cortex of the mouse are specialized in the processing of specific acoustic features present in human speech. These results complement extensive literature on speech processing in other animals, including rat, cat, and primate (Porter et al., 2011; Centanni et al., 2013; Engineer et al., 2015; Eggermont, 1995; Wong and Schreiner, 2003; Steinschneider et al., 2003; Tsunada et al., 2011), and enable the use of advanced tools readily available in the mouse to dissect the neural circuits responsible for speech processing. While acoustic communication in humans involves speech-production anatomical features and complex grammatical structures that are not expected to be well modeled in the mouse, the results above provide evidence that the auditory system of the mouse can effectively process acoustic features of human speech, allowing for a detailed investigation of the neural mechanisms underlying speech processing and speech-sound learning.

A potential limitation of studying speech sound processing using the mouse is the clear difference in frequency hearing range between humans and mice. To address this difference, human speech sounds were shifted up in frequency to the mouse hearing range. The assumption here is that the features of speech under study can be maintained while varying other sound features. While this adjustment can be easily performed for the perception of spectral features, it is more challenging to identify appropriate adjustments for temporal features. For instance, our evaluation of categorization performance using different sets of formant transition slopes yielded varied results, with mice learning the task better when using slopes closer to those naturally occurring in their vocalizations, compared to those common in human speech. This finding further emphasizes the importance, when conducting animal studies of human speech processing, of selecting sounds that contain the features of interest within an ethologically relevant range given a particular experimental animal model.

In studies with human subjects, functional imaging and neurophysiological investigations have revealed the existence of distinct cortical regions responsible for processing various speech attributes, such as phonemic information, prosody, and voice characteristics (Yi et al., 2019). Therefore, for an animal model to be valid for the study of speech perception, it would be expected that distinct neural circuits specialize in the processing of different features of speech. Consistent with this idea, we found that the dorso-posterior region of the auditory cortex of the mouse contains a higher proportion of neurons that are selective to the speech features tested, compared to other cortical regions.

Differential roles across neural circuits, however, were not readily apparent when we grouped neurons according to atlas-defined cortical areas. Several possibilities could explain these discrepancies. First, a technical limitation of our study is that brain areas were defined by registering each mouse brain to the atlas, yet it is known that the extent of these areas (when functionally defined) can vary significantly across animals (Narayanan et al., 2023). Consequently, the assignment of each neuron to a given brain area may lack precision, potentially affecting the interpretation of comparisons across atlas-defined areas. Future studies where brain areas are functionally defined for each animal (*e.g.*, via widefield imaging of tonotopic organization) could ameliorate this issue. A second interpretation of the lack of differences across the cortical areas measured is that only subregions within these brain areas could be specialized in processing each feature. Therefore, a higher level of granularity in how areas are parcellated beyond standard mouse brain atlases we used would be needed to identify specializations within the auditory cortex. In fact, various ways of parcellating the auditory cortex of the mouse have been proposed, ranging from a “lumper” approach yielding only a few areas to a “splitter” approach with many more areas (Ceballo et al., 2019; Issa et al., 2017; Tsukano et al., 2016; Romero et al., 2020). Implementing alternative parcellation strategies may provide a more detailed understanding of functional specialization of speech feature processing within the auditory cortex of the mouse.

In summary, while animal models are not expected to capture all the complexities of human speech perception, our study provides support for the mouse as a useful model for investigating the neural circuits that underlie the processing the basic acoustic features of speech and how these circuits change during learning new sound contrasts, as it occurs during second language acquisition.

## Acknowledgements

We thank Gabriel Toea and Diyar Dezay for assistance with animal training.

## References

1. Ceballo S, Piwkowska Z, Bourg J, Daret A, Bathellier B (2019) Targeted Cortical Manipulation of Auditory Perception. Neuron 104:1168–1179.e5.

2. Centanni TM, Engineer CT, Kilgard MP (2013) Cortical speech-evoked response patterns in multiple auditory fields are correlated with behavioral discrimination ability. Journal of Neurophysiology 110:177–189.

3. Eggermont JJ (1995) Representation of a voice onset time continuum in primary auditory cortex of the cat. The Journal of the Acoustical Society of America 98:911–920.

4. Engineer CT, Rahebi KC, Buell EP, Fink MK, Kilgard MP (2015) Speech training alters consonant and vowel responses in multiple auditory cortex fields. Behavioural Brain Research 287:256–264.

5. Hammerschmidt K, Reisinger E, Westekemper K, Ehrenreich L, Strenzke N, Fischer J (2012) Mice do not require auditory input for the normal development of their ultrasonic vocalizations. BMC Neuroscience 13:40.

6. Issa JB, Haeffele BD, Young ED, Yue DT (2017) Multiscale mapping of frequency sweep rate in mouse auditory cortex. Hearing Research 344:207–222.

7. Jadoul Y, Thompson B, De Boer B (2018) Introducing Parselmouth: A Python interface to Praat. Journal of Phonetics 71:1–15.

8. Kluender KR (2000) Contributions of nonhuman animal models to understanding human speech perception. The Journal of the Acoustical Society of America 107:2835–2835.

9. Kuhl PK, Miller JD (1978) Speech perception by the chinchilla: Identification functions for synthetic VOT stimuli. The Journal of the Acoustical Society of America 63:905–917.

10. Kuhl PK, Padden DM (1983) Enhanced discriminability at the phonetic boundaries for the place feature in macaques. The Journal of the Acoustical Society of America 73:1003–1010.

11. Lotto A, Kluender K, Holt L (2003) Animal Models of Speech Perception Phenomena. Chicago Linguist. Soc. 33.

12. Mahrt EJ, Perkel DJ, Tong L, Rubel EW, Portfors CV (2013) Engineered Deafness Reveals That Mouse Courtship Vocalizations Do Not Require Auditory Experience. The Journal of Neuroscience 33:5573–5583.

13. Narayanan DP, Tsukano H, Kline AM, Onodera K, Kato HK (2023) Biological constraints on stereotaxic targeting of functionally-defined cortical areas. Cerebral Cortex 33:3293–3310.

14. Oganian Y, Bhaya-Grossman I, Johnson K, Chang EF (2023) Vowel and formant representation in the human auditory speech cortex. Neuron 111:2105–2118.e4.

15. O’Sullivan C, Weible AP, Wehr M (2020) Disruption of Early or Late Epochs of Auditory Cortical Activity Impairs Speech Discrimination in Mice. Frontiers in Neuroscience 13:1394.

16. Pachitariu M, Steinmetz N, Kadir S, Carandini M, Harris KD (2016) Kilosort: realtime spike-sorting for extracellular electrophysiology with hundreds of channels. bioRxiv p. 061481.

17. Porter BA, Rosenthal TR, Ranasinghe KG, Kilgard MP (2011) Discrimination of brief speech sounds is impaired in rats with auditory cortex lesions. Behavioural Brain Research 219:68–74.

18. Portfors CV (2007) Types and Functions of Ultrasonic Vocalizations in Laboratory Rats and Mice. Journal of the American Association for Laboratory Animal Science 46.

19. Portfors CV, Perkel DJ (2014) The role of ultrasonic vocalizations in mouse communication. Current Opinion in Neurobiology 28:115–120.

20. Romero S, Hight AE, Clayton KK, Resnik J, Williamson RS, Hancock KE, Polley DB (2020) Cellular and Widefield Imaging of Sound Frequency Organization in Primary and Higher Order Fields of the Mouse Auditory Cortex. Cerebral Cortex 30:1603–1622.

21. Rossant C, Kadir SN, Goodman DFM, Schulman J, Hunter MLD, Saleem AB, Grosmark A, Belluscio M, Denfield GH, Ecker AS, Tolias AS, Solomon S, Buzsáki G, Carandini M, Harris KD (2016) Spike sorting for large, dense electrode arrays. Nature Neuroscience 19:634–641.

22. Saunders JL, Wehr M (2019) Mice can learn phonetic categories. The Journal of the Acoustical Society of America 145:1168–1177.

23. Steinschneider M, Fishman YI, Arezzo JC (2003) Representation of the voice onset time (VOT) speech parameter in population responses within primary auditory cortex of the awake monkey. The Journal of the Acoustical Society of America 114:307–321.

24. Tsukano H, Horie M, Hishida R, Takahashi K, Takebayashi H, Shibuki K (2016) Quantitative map of multiple auditory cortical regions with a stereotaxic fine-scale atlas of the mouse brain. Scientific Reports 6:22315.

25. Tsunada J, Lee JH, Cohen YE (2011) Representation of speech categories in the primate auditory cortex. Journal of Neurophysiology 105:2634–2646.

26. Wang Q, Ding SL, Li Y, Royall J, Feng D, Lesnar P, Graddis N, Naeemi M, Facer B, Ho A, Dolbeare T, Blanchard B, Dee N, Wakeman W, Hirokawa KE, Szafer A, Sunkin SM, Oh SW, Bernard A, Phillips JW, Hawrylycz M, Koch C, Zeng H, Harris JA, Ng L (2020) The Allen Mouse Brain Common Coordinate Framework: A 3D Reference Atlas. Cell 181:936–953.e20.

27. Wong SW, Schreiner CE (2003) Representation of CV-sounds in cat primary auditory cortex: intensity dependence. Speech Communication 41:93–106.

28. Yi HG, Chandrasekaran B, Nourski KV, Rhone AE, Schuerman WL, Howard MA, Chang EF, Leonard MK (2021) Learning nonnative speech sounds changes local encoding in the adult human cortex. Proceedings of the National Academy of Sciences 118:e2101777118.

29. Yi HG, Leonard MK, Chang EF (2019) The Encoding of Speech Sounds in the Superior Temporal Gyrus. Neuron 102:1096–1110.

